## Supplementary Information for "Non-canonical function of folate/folate receptor 1 during neural tube formation"

**Supplementary Table 1. FOLR1-interacting proteins during neural plate stages.**
Lysates from neural plate stage *Xenopus laevis* embryos were immunoprecipitated with anti-FOLR1 antibody and immunoprecipitates (IP) were processed for LC-MS/MS assays. Shown are numbers of peptides identified for a selection of FOLR1-interacting proteins in each run.

### Supplementary Fig. 1

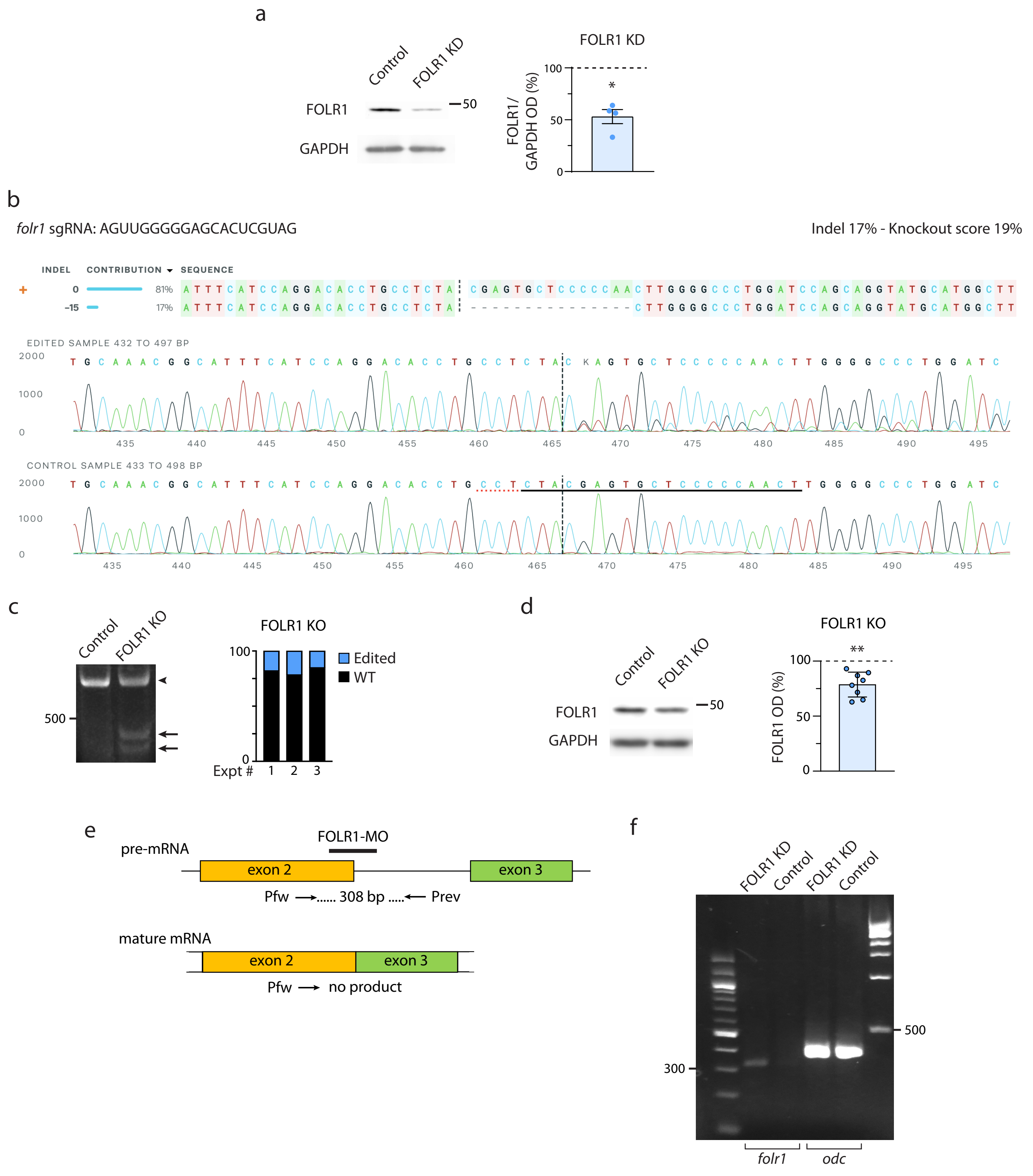

### Supplementary Fig. 2

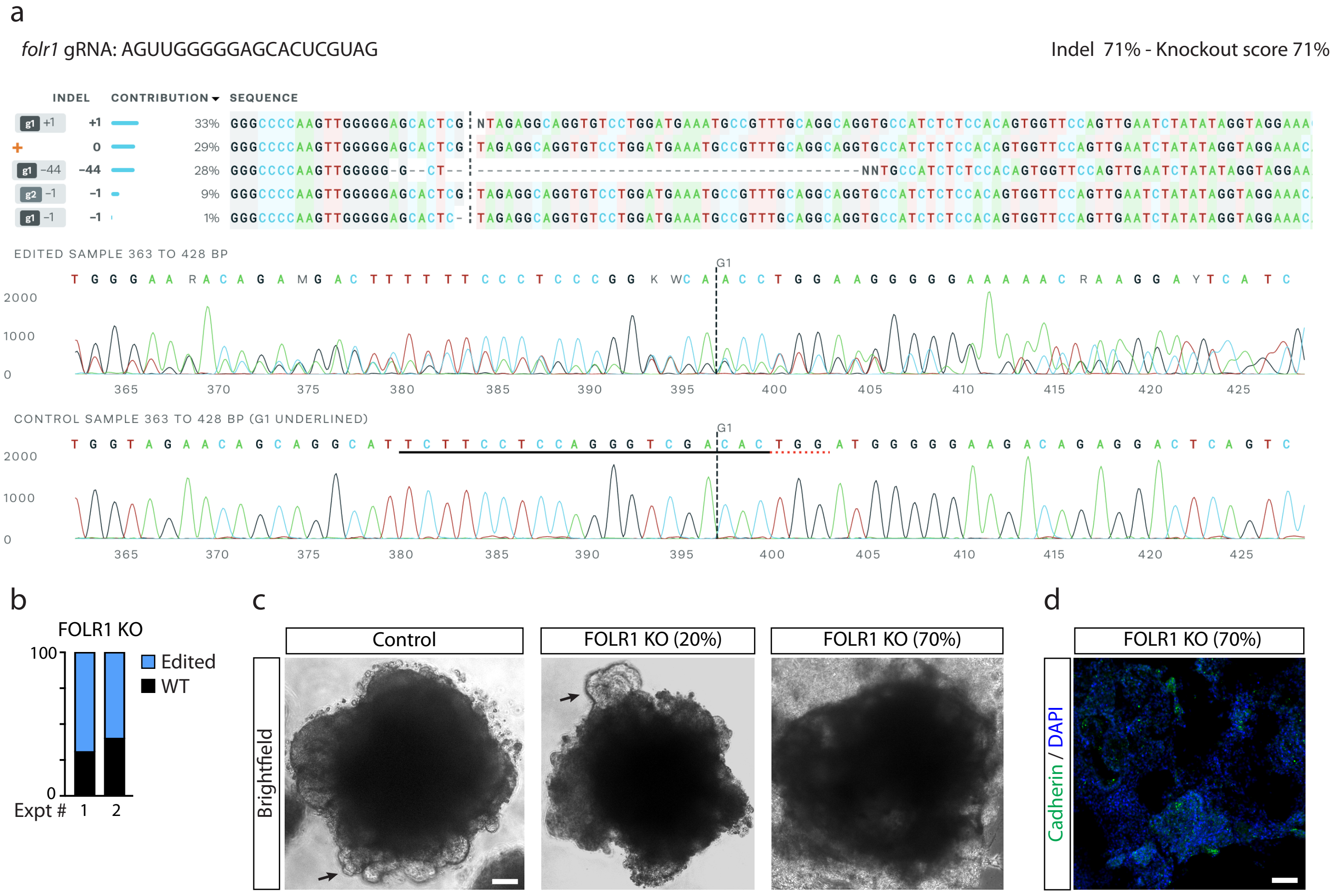

Supplementary Fig. 3

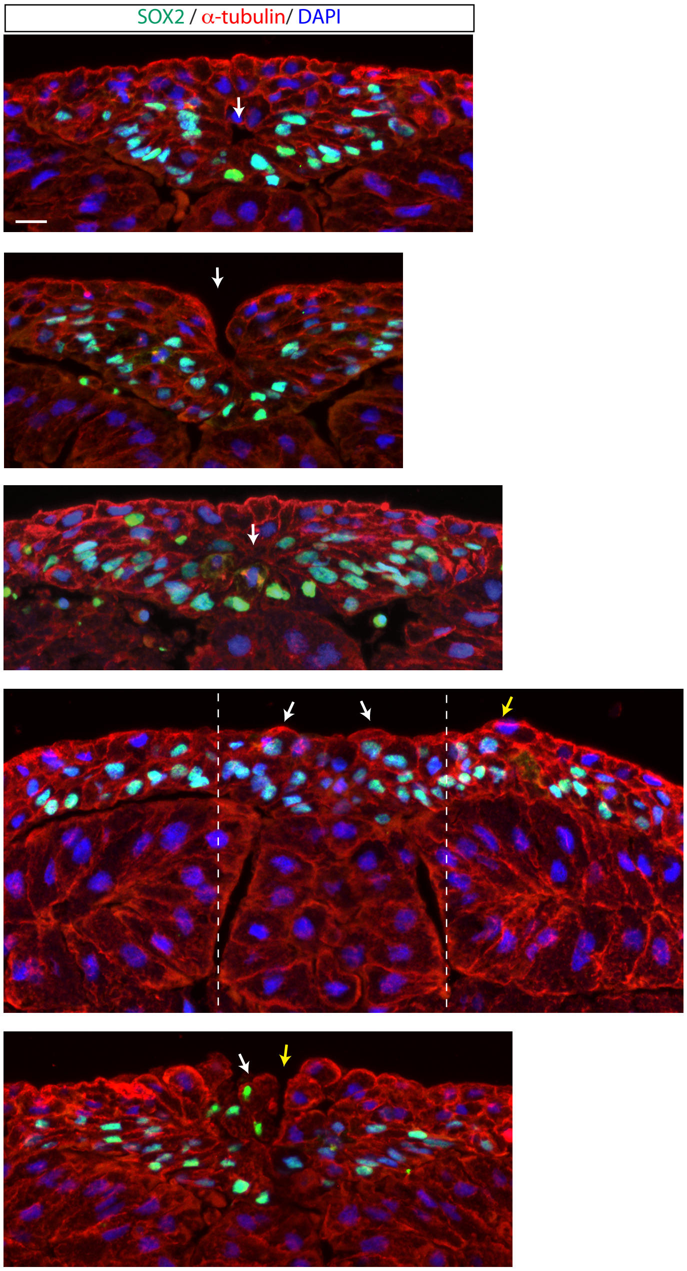


| Neural tissue defect score |
|---|
| 0 |
| 1<br>Open neural tube: 1 |
| 1<br>No lumen: 1 |
| 3<br>Open neural tube: 1<br>Unfolded neural plate: 1<br>Rounded cells: 1 |
| 3<br>Open neural tube : 1<br>Neural tissue disruption: 1<br>Rounded cells: 1 |

Supplementary Fig. 4

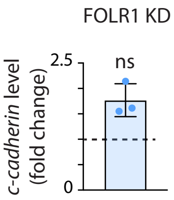

Supplementary Fig. 5

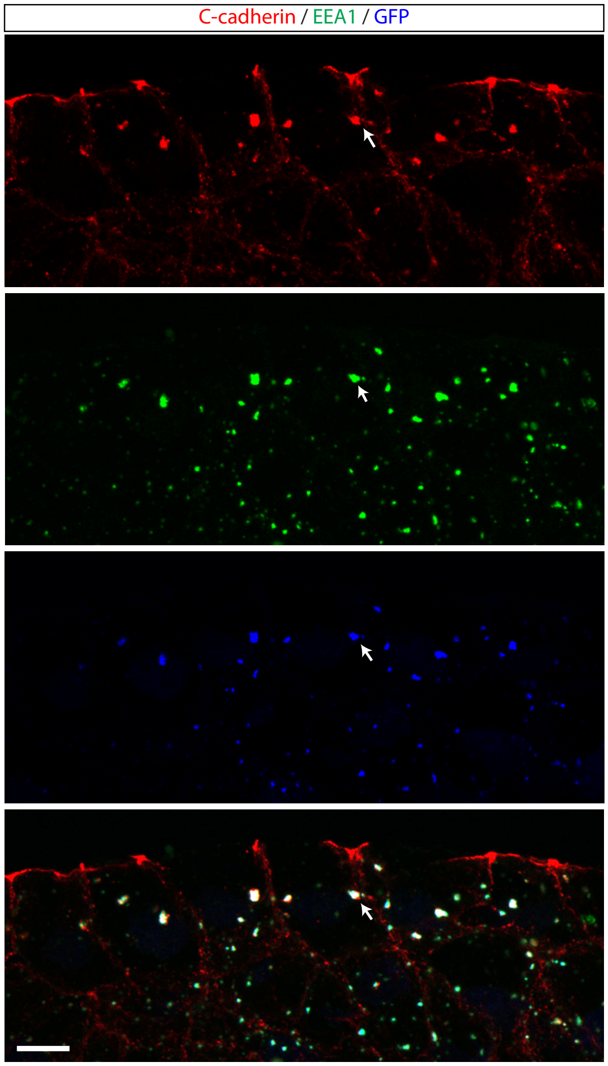

Supplementary Fig. 6

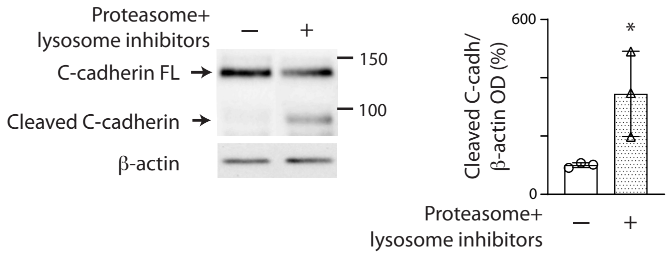

Supplementary Fig. 7

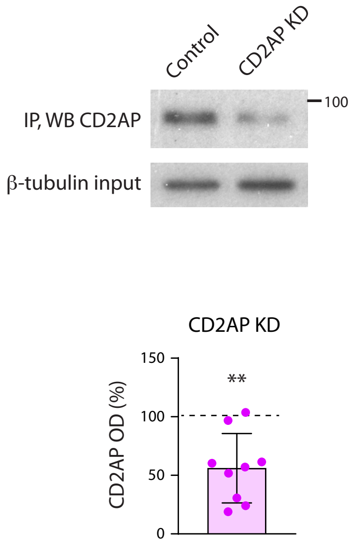

**Supplementary Table 1**

| Protein name/ UniProt accession # | Replicate | # Peptides identified FOLR1-IP | # Peptides identified Control-IP |
|---|---|---|---|
| **β-catenin Ctnnb1/** A1A5I6 | 1 | 4 | 0 |
| | 2 | 5 | 0 |
| | 3 | 4 | 0 |
| **C-cadherin Xb-cadherin/** Q6NTM0 | 1 | 1 | 0 |
| | 2 | 7 | 0 |
| | 3 | 4 | 0 |
| **CD2AP/** Q4V7X8 | 1 | 2 | 0 |
| | 2 | 2 | 0 |
| | 3 | 2 | 0 |
